# Harnessing within-cultivar variation to identify hidden genetic resistance using single plant-omics

**DOI:** 10.64898/2026.04.02.710456

**Authors:** Ethan J Redmond, Meng Li, Samuel Holden, Muhammad Jawad Akbar Awan, Yishan Zhang, Jujhar Singh Gill, Jashanpreet Virhia, Jess Hargreaves, Peter Danks, Beth Sleath, Rajagopal Subramaniam, Carmen Hicks, David P Overy, Gurcharn Singh Brar, Daphne Ezer

**Affiliations:** Department of Biology, University of York, Heslington, York United Kingdom, YO10 5DD; Faculty of Agricultural, Life and Environmental Systems, University of Alberta, Edmonton, AB, Canada, T6G 2R3; Faculty of Land and Food Systems, The University of British Columbia, Vancouver, BC, Canada, V6T 1Z4; Department of Mathematics, University of York, Heslington, York United Kingdom, YO10 5DD; Agriculture and Agrifood Canada Ottawa Research and Development Centre, Ottawa, ON, Canada, K1A 0C6

## Abstract

*Fusarium graminearum* is a fungal pathogen that causes scab or head blight in small grain cereals and threatens global cereal production. Disease progression varies widely among individual plants of the same cultivar, reflecting both genetic and environmental heterogeneity. This complicates the identification of early host responses, because each individual plant in a population is at a slightly different phase of disease progression. Here we apply single plant-transcriptomics to a population of 121 malt-barley exposed to *F. graminearum*, enabling us to reconstruct a temporal sequence of regulatory processes during early infection. We identified several disease-resistance associated genetic variants that are already endemic in this high-yielding cultivar, suggesting potential as breeding targets. These variants were within proteins involved in ROS-burst production, a lectin-kinase PRR, and enzymes with DON-detoxification activity. Single plant-transcriptomics offers a novel strategy for characterising early plant-pathogen interactions, turning intra-population heterogeneity from an experimental barrier into an asset.

---

Fusarium head blight (FHB) or scab, caused by the fungal pathogen *Fusarium graminearum* Schwabe, is a seed and residue-borne disease of cereal crops such as wheat, barley, oat, and rye. Currently the disease is widespread in North America, and around the Mediterranean sea, but it is thought to have some presence in all wheat growing countries in the world^1,2,3^.

*Fusarium graminearum* infection causes direct yield losses through infecting plant vascular tissue (wilt), and at anthesis through penetration of the developing seed and subsequent necrotrophic growth (head blight) which can cause total loss of yield in favourable conditions. A serious problem associated with FHB infection is the production of mycotoxins in the grain leading to grade loss. Of the various different mycotoxins associated with FHB in barley, deoxynivalenol (DON) produced by *F. graminearum* is of a primary concern as it affects malting quality and can cause vomiting or intestinal distress when ingested by humans or used as animal feed^4,5^. In some cases a fungal infection will produce enough DON to render the seed unusable, even though the yield appears high^6^. The Canadian Grain Commission’s survey estimated economic losses due to FHB predicted to be 1 billion CAD in 2016.

A challenge in understanding how plants respond to *F. graminearum* infection is the heterogeneity of the plant’s response, even within a single cultivar^7,8^. This contrasts with many other cereal diseases such as rusts or powdery mildews where resistant crops tend to be very consistent in their phenotypes^9^. In barley, heterogeneity in response to *F. graminearum* infection is derived from both intrinsic and extrinsic sources. Intrinsic heterogeneity stems from genetic factors governing the response to infection; for instance, genetic factors associated with awn physiology often impact *Fusarium* resistance in barley^10^. Extrinsic factors include the density and viability of the initial spore inoculum, as well as microenvironmental conditions influencing spore germination and heterogeneous infection rates. Furthermore, *F. graminearum* strains vary in DON isoform production and secretion rates, that further contribute to variable disease symptoms^11,12^. Consequently, at any given time post infection, individual plants will be at different phases of disease progression and/or immune response.

Single plant-omics provides a framework to resolve these asynchronous biological processes. In single cell RNA-seq analysis, individual cells are ordered by their transcriptional trajectories to produce a pseudo-time series, often revealing developmental or temporal patterns in expression^13^. Similarly, performing transcriptomics on a large population of individual plants permits *post-hoc* ordering by transcriptional trajectories, in our case ordering the barley plants based on transcriptional signatures of infection progression. We can then map how physiological factors and gene expression values change over the pseudo-time course, to determine the order of transcriptional events that occur during the infection process and identify specific alleles or gene expression variation patterns that may contribute to infection progression. Single plant-omics has previously been used to dissect transcriptomic variation associated with yield heterogeneity in field trials^14^, describe gene expression noise over a diurnal cycle^15^, and unravel the transcriptional dynamics during developmental phase transitions^13,16^. However, this technique has not previously been deployed to investigate immune responses in any plant species.

Additionally, we identified genetic variants contributing to the heterogeneity in *F. graminearum* responses in the cultivar, as no breeding process completely eliminates SNPs from the seed pool. Traditionally, variants associated with disease response are identified through a genome-wide association study (GWAS) across genetically diverse lines^17^. Here, we show that it is possible to identify significant variants associated with disease traits using single plant-transcriptomics. We also show that these variants are associated with disease-related traits in a validation set of plants (not used in the initial GWAS analysis), demonstrating that this phenomenon is not a consequence of overfitting. Moreover, many of the variants are in genes that are known to be associated with disease-related traits, suggesting causative links.

In conclusion, we leverage recent innovations in single plant-omics to identify genes that are differentially regulated during early *F. graminearum* infection in barley. Several promising genetic variants within an elite cultivar that are significantly associated with disease progression were discovered. This study demonstrates that single plant-omics is a promising framework for resolving asynchronous dynamics of disease progression in cultivated crops.

## Results

### Differentially expressed genes in *F. graminearum*-treated barley are associated with disease severity in individual plants

A single plant-transcriptomics experiment was performed on cv. AAC Synergy barley plants (**Fig. 1**), derived from three self-pollinated plants from the breeder seed source. AAC Synergy is an elite, popular Canadian malt barley cultivar, sharing alleles with many other popular field cultivars^18^. It has intermediate FHB resistance and heterogeneous responses between individuals, making it well suited to capture a broad range of disease responses^18^.

**Figure 1.**
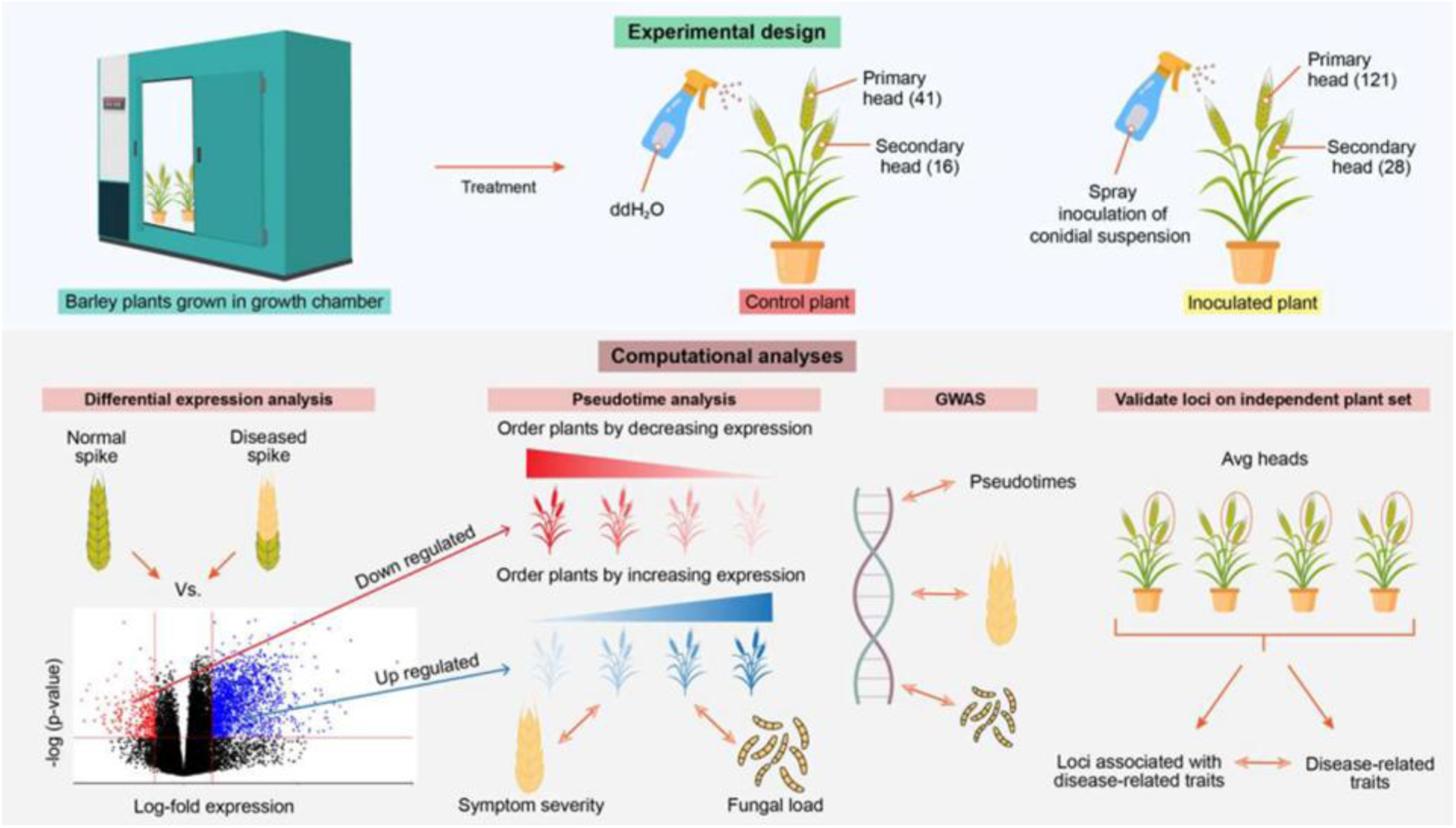
Workflow for using single plant-omics for identifying early markers of Fusarium infection in barley.

A total of 121 plants were spray inoculated with *F. graminearum* spores (*Fg*-treated) and 41 were mock-treated. For each sample, we measured phenotypic factors including plant height, Fusarium Damaged Kernels (FDK) and total kernels at three timepoints post-inoculation, final head weight, and DON concentration in head tissue (**Table S1**). RNA-seq was performed using the topmost two spikelets harvested five days after inoculation, to measure gene expression and identify genetic variants in the coding sequences (**Table S2**). As breeder seed source was used, the line was extremely pure. RNA-seq identified a total of 3910 SNPs in exons across the population relative to the expected reference, although it is important to note that variant calling quality is correlated to expression levels and so there are likely many undiscovered SNPs in non-coding or poorly expressed genes.

Using a Principal Component Analysis (PCA) of genetic variants, we found that there were two sub-populations of barley in our sample (**Fig. S1A**), driven by genetic variants in unplaced scaffolded genomic regions, rather than in the assembled chromosomes (**Fig. S1B**). In addition to excluding lowly expressed genes (**Fig. S1C**) in further analysis, we excluded genes within scaffolds and those that were differentially expressed between the two sub-populations, so the impact of the sub-population structure does not dominate our results (**Fig. S1C**). The treatment and control plants were indistinguishable in the PCA of genetic variants, confirming that the genotypic patterns in the data were not primarily driven by infection-associated expression patterns (**Fig. S1A**).

Next, genes were identified that were differentially expressed between treatment and control plants, as well as genes that differed between the top and bottom quartiles of disease severity, as indicated by the Area Under the Disease Progression Curve (AUDPC) (**Table S3**). The RNA-seq was performed after 5 days of infection, while disease traits were measured after 20 days, so these differentially expressed gene sets are indicative of early infection responses that predict future disease outcomes.

Plants with more severe symptoms (larger AUDPC) showed increased expression of heme binding and monooxygenase activity (**Fig. 2A**), while lipid metabolic process genes were more expressed in plants with smaller AUDPC (**Fig. S2A**). Genes involved in heme binding and monooxygenase activity included Cytochrome P450 (CYP450s) which are critical for defense in plants via detoxification^19,20^. Genes more highly expressed both in *Fg*-treated plants and in plants with larger AUDPC included those related to tetrapyrrole binding and oxoacid metabolic processes (**Fig. S2B**), perhaps suggesting pathogen-triggered metabolic reprogramming^21,22^.

**Figure 2.**
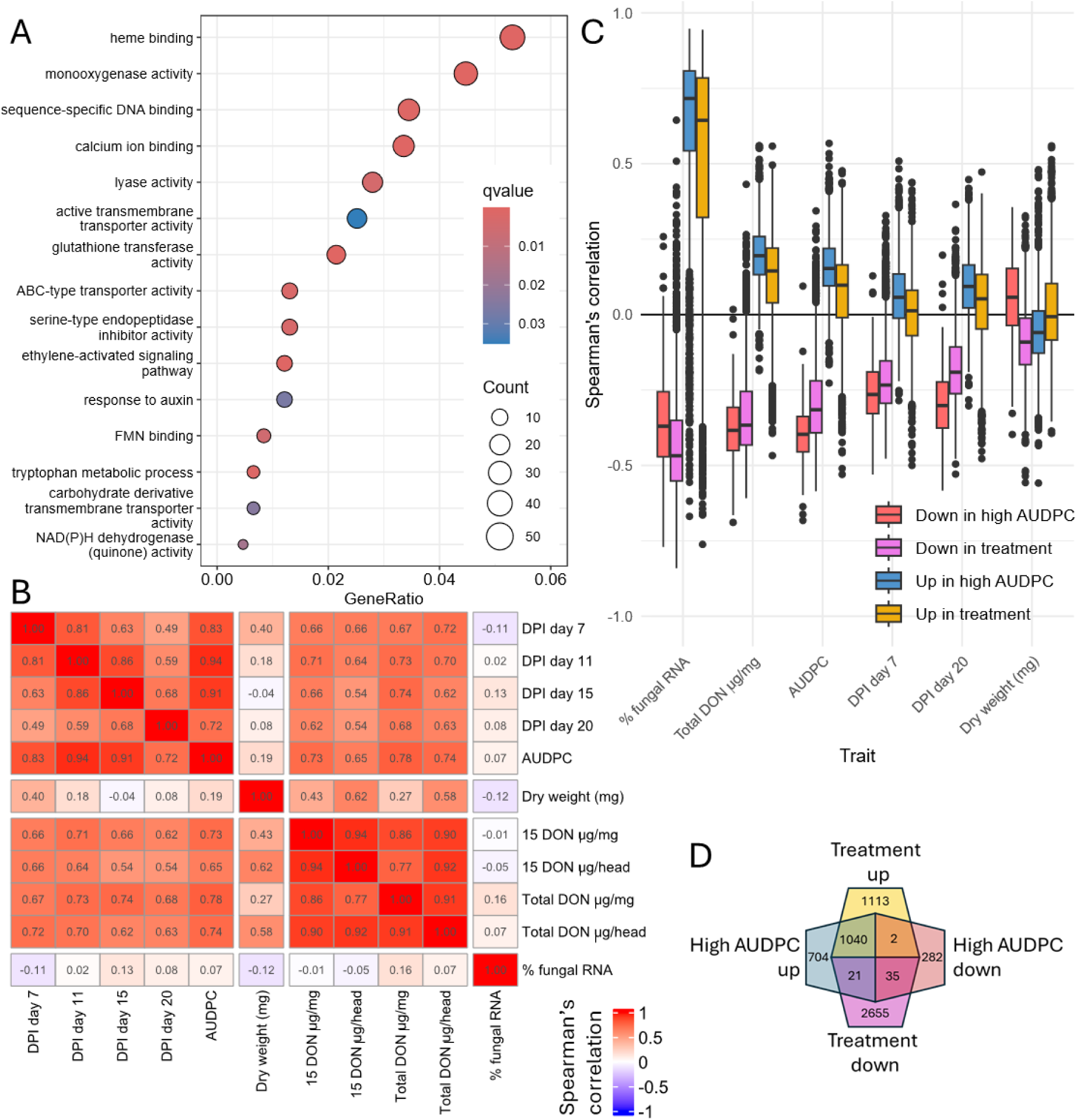
Differentially expressed genes under Fusarium infection. Panel (A) shows representative Gene Ontology (GO) terms associated with genes that were more highly expressed in the top 25% Area Under the Disease Progressive Curve (AUDPC) plants than the bottom 25%. Panel (B) shows the Spearman’s correlation between dry weight, fungal load (% fungal RNA), and disease-related traits, specifically the disease progression index (DPI) on days 7-20, AUDPC, and alternative measures of DON toxin concentrations. Panel (C) shows the distribution of Spearman’s correlations between gene expression values and each trait shown in (B), calculated separately for each set of differentially expressed genes (genes whose expressions go up and down in the plants with the top vs bottom 25% AUDPC and those whose expressions go up or down in treatment vs control barley plants). Boxplots indicate the median, interquartile ranges, and range, with outliers lying >1.5*IQR away from the median. Panel (D) shows the overlap between these gene sets, with numbers indicating gene counts.

The proportion of fungal transcripts in the RNA-seq samples can be used to estimate the fungal load in the spikelets for each individual plant. Although AUDPC and DON-related traits were positively correlated with each other, percent fungal RNA was not strongly associated with any disease severity trait (**Fig. 2B**). Genes that were more expressed in *Fg-*treatment were more strongly associated with percent fungal RNA than other disease-related traits. Intriguingly, genes with *reduced* expression under *Fg-*treatment were more strongly associated with AUDPC than genes with increased expression (**Fig. 2C**). The same pattern emerged when comparing genes that were differentially expressed between the top quartile vs. bottom quartile AUDPCs. These results suggest strong early infection response may curtail symptom severity, but early down-regulation of non-stress/infection associated processes may generate more severe future symptoms.

### A transcriptional trajectory reveals defence processes associated with fungal transcript abundance

Transcriptional profiles of *F. graminearum*-infected barley revealed substantial expression heterogeneity between plants (**Fig. S3**); notably, these gene expression levels correlated significantly with disease-associated traits (**Fig. 2C**). These data indicate an underlying transcriptional trajectory (pseudotime) among infected barley plants, reflecting the asynchronous progression of the disease. Peudotime was defined as an ordering of individual plants in which genes in a specified set change their expression monotonically, as in Redmond et al^13^. Two separate pseudotime rankings were constructed on a training set of *F. graminearum* infected plants (n=93), using the sets of genes that showed either higher or lower expression in *Fg*-treated plants compared with controls.

Pseudotimes were found to be highly robust across 300 random gene-set partitions (**Fig. S4AB**) and reflected the positioning of plants across the first principal component of a PCA of gene expression (**Fig. S4CD**). The genes that were more highly expressed in *Fg*-treated plants formed several clusters whose expression varied over pseudotime (**Fig. 3A**). There were three large clusters that increased expression over pseudotime, that were designated as early, medium, and late based on the order in which gene expression began to increase over the pseudotime. A representative selection of gene ontology (GO) terms for these clusters are shown (**Fig. 3B**, **Table S4**), suggesting that processes such as UDP-glycosultransferase, calcium ion binding and chitinase activity (early) may be up-regulated sooner in the infection timeline than hormone-mediated signalling and xenobiotic transport (medium). Transcripts associated with the extracellular matrix and strictosidine synthase were associated with late expression. In contrast, the pseudotime generated using genes with reduced expression under *F*.

**Figure 3.**
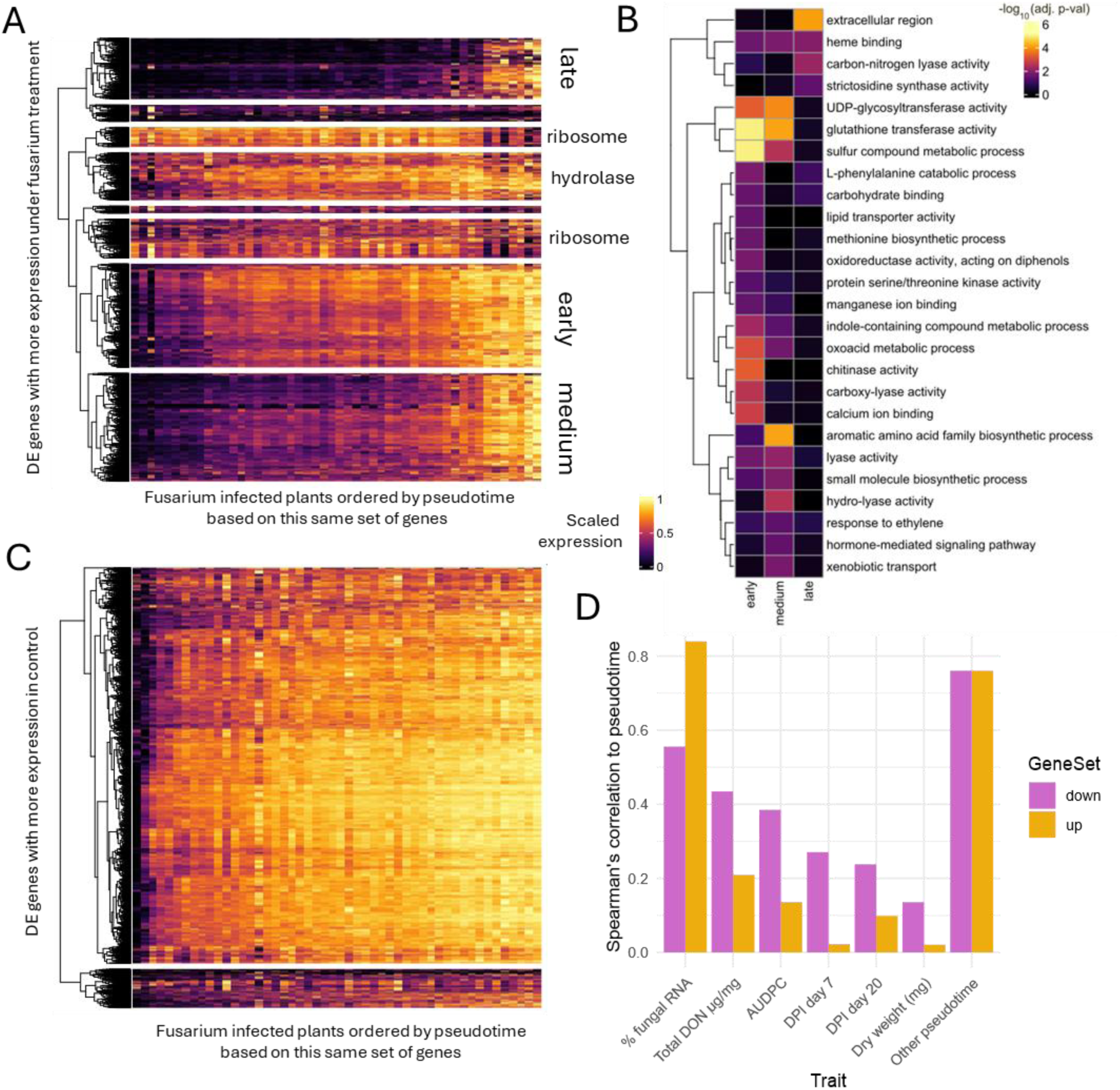
Transcriptional trajectory among genes that go up and down under Fusarium treatment. First, a pseudotime was inferred using differentially expressed genes whose expression was higher under fusarium treatment than mock-treated plants. Panel (A) shows the expression of these genes over pseduotime, with clusters determined by complete linkage, and expression patterns scaled between 0 (minimum expression) and 1 (maximum expression). GO enrichment analysis was performed on each cluster of genes, with two clusters showing primary enrichment for ribosomes, one cluster showing only enrichment for hydrolases, and the three remaining large clusters (labelled early, medium, and late) showing enrichment for various GO terms, with representative examples shown in (B). In (C), the same analysis as in (A) was performed using only genes that had higher expression under the treatment, but this showed fewer distinct clusters. (D) highlights the Spearman’s correlation between each pseudotime and associated traits.

*graminearum* infection did not produce a strong clustering of gene expression values (**Fig. 3C**), which suggests that gene repression may be less tightly coordinated than gene activation during response to infection. While specific genes are likely to be switched off via effectors, these genes will not be present in sufficient numbers to be detected by GO-term analysis relative to the overall gene-space.

The sequence of gene expression events in response to *F. graminearum* challenge is intuitive. Early stage genes (calcium ion signalling, general-purpose chitinase, and UDP-glycosultransferase for DON detoxification) are all responses the plant immediately takes as part of sensing and responding to pathogen presence^23,24,25^. Meanwhile, hormone signalling (in this case jasmonic acid and strictosidine pathways) is a downstream effect of defence signalling^26,27^, as individual cells switch from growth to defence and communicate across the plant tissue.

Each plant would be expected to be at different phases of disease progression due to different fungal loadings. We hypothesized that the proposed plant orderings we developed (pseudotimes) represent the progression of response to *F.* graminearum infection. Supporting this hypothesis, we found that both pseudotimes (developed using either up-or down-regulated genes) were more strongly correlated with disease symptom-related traits than dry weight (**Fig. 3D**). The pseudotime associated with up-regulated genes was more strongly associated with percent fungal RNA than the one made using down-regulated genes, but the inverse was true of AUDPC and DON, consistent with our earlier findings.

### Genetic variants in an elite barley cultivar are associated with disease progression

Despite being a single cultivar, AAC Synergy harbours some genetic heterogeneity. We hypothesized that some of these variants may be associated with disease symptom severity, which could serve as potential markers for breeding for *Fusarium* resistance. Moreover, as there are sparse numbers of gene variants within a single cultivar (with all variants shown in **Fig. S5**), we hypothesized that GWAS conducted on this dataset may be enriched for causal variants compared to a traditional GWAS, where the causal variants are often hidden within large linkage disequilibrium blocks.

Supporting this hypothesis, we performed a GWAS and found that nearly all the variants associated with traits of interest lie within genes whose functions have previously been linked with the plant immune regulatory network (**Fig. 4, Table S6**). Variants associated with pseudotime were found to be associated with ROS production^28^, gylycosyltransferase activity^25^, receptor kinase activity and downstream CDPK activity^29^, and transcriptional reprogramming (**Fig. S6**). Over 60% of variants that were found to be associated with pseudotimes and disease phenotypes were within gene bodies of known plant immunity genes. These results suggest that performing GWAS within a single near isogenic population may be an effective strategy for identifying intra-population variants associated with traits of interest that may be causal.

**Figure 4.**
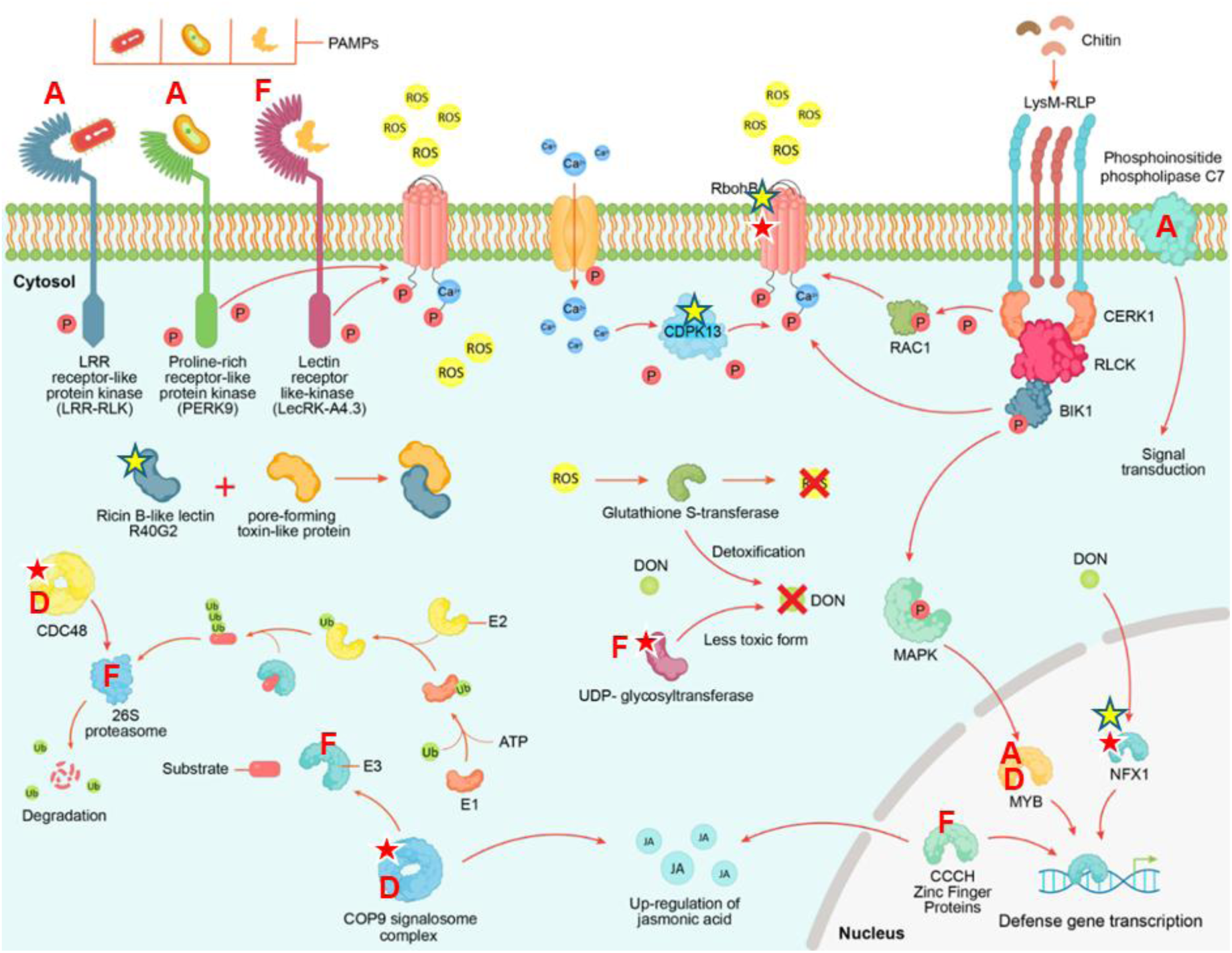
Proposed mechanism of how endemic genetic heterogeneity in AAC Synergy contributes to variation in Fusarium head blight progression. Here we summarise the previously known fusarium response pathways in cereals, overlaying symbols to indicate which proteins have endemic SNPs associated with disease traits in AAC Synergy based on a Genome-Wide Association Study (GWAS). Note that over 60% of all variants that were found to be associated with disease traits are shown in this figure, so most variants are directly within the gene bodies of known components of plant immunity. Yellow stars indicate variants within genes that are associated with the pseudotime made from genes up-regulated during infection and the red stars are for variants associated with the pseudotime made from genes that were down-regulated during infection. Red letters designate variants within genes associated with other key disease-related traits: A for AUDPC, D for total DON concentration, and F for percent fungal RNA in the sample. All other named genes within this figure were found to be differentially expressed between treatment and control barley plants.

### Genetic variants can be used as markers of disease-related traits

Next, we assessed whether the identified genetic variants would indicate *F. graminearum* disease traits in an independent set of barley plants that were not used in the pseudotime or GWAS analysis (n=28) collected in parallel with the initial experiment.

One challenge is that each barley spike experiences a different fungal load due to the spray-based inoculation strategy, which is further exacerbated by temporal variation in *F. graminearum* spore germination and infection events. These factors increase heterogeneity in disease symptoms due to factors that are independent of the plant’s genetic background. To reduce this environment-driven variability, we include two technical replicates, i.e., samples taken from two heads of the same barley plant. These two heads share the same genetic background but are exposed to distinct fungal loads. Disease-related traits were positively associated with each other across technical replicates (**Fig. 5A**). To reduce the heterogeneity in these traits caused by environmental factors, we took an average across both replicates in subsequent analysis. 17 genetic variants that were found in at least one GWAS were present in our validation set (**Fig. S7**).

**Figure 5.**
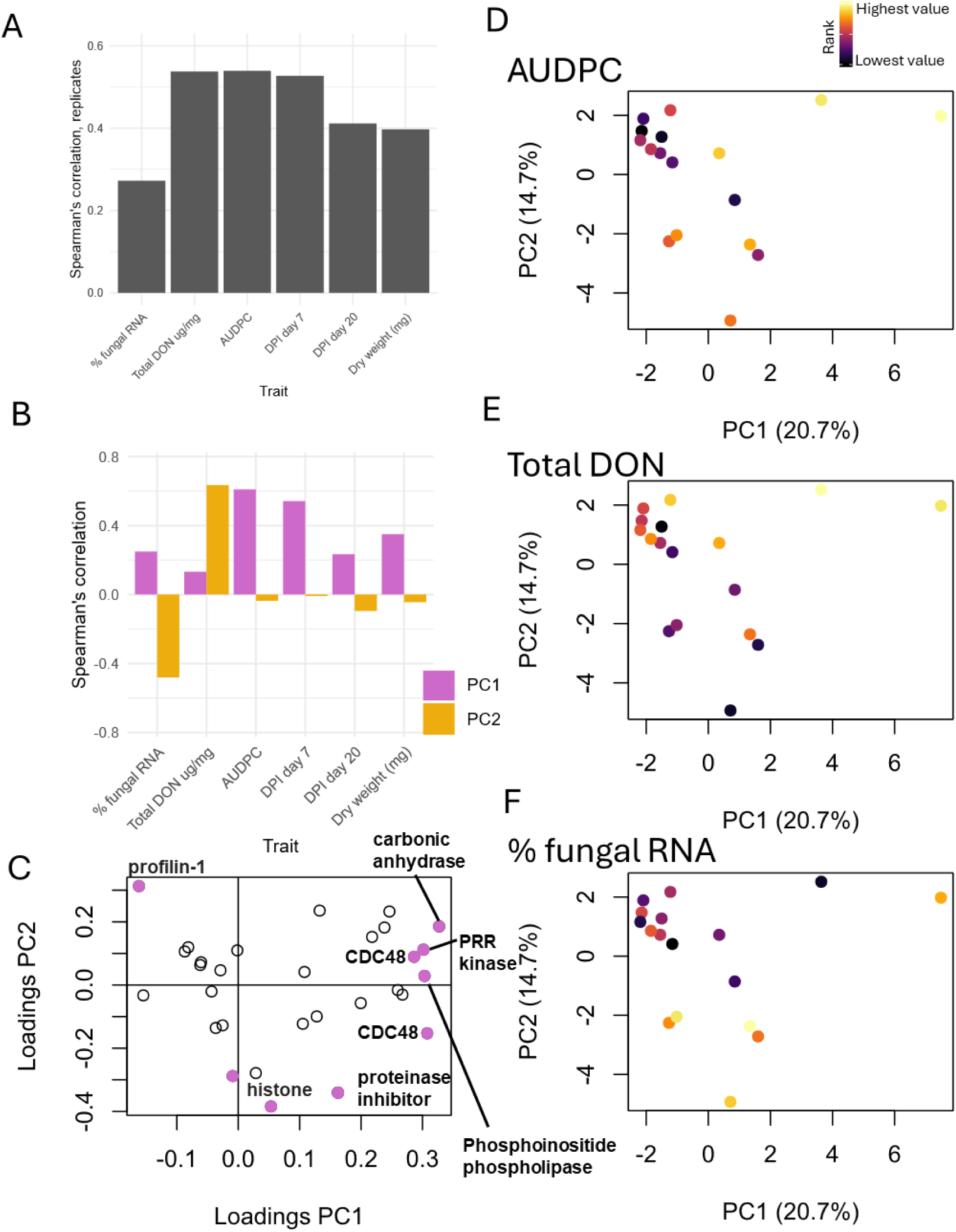
Variants can be used to predict disease-related traits. (A) Spearman’s correlation between traits across pairs of heads. (B) Spearman’s correlations between principal components and traits. (C) Loadings of principal components (PCs), with several genes with high values highlighted. PRR kinase refers to profile-rich receptor-like kinase and CDC48 refers to a cell division cycle protein 48. The ranks of (D) the Area under the disease progression curve (AUDPC) (E) Total DON toxin concentration and (F) % fungal RNA are used to colour-code a principal component analysis (PCA) plot, with percentage of variance explained by each PC indicated in brackets.

We confirmed that these variants were associated with disease symptoms. PC1 was more highly correlated with AUDPC, especially how large the lesion was in the earliest time point, Disease Progression Index (DPI) day 7, suggesting that PC1 captures how quickly lesions form (**Fig. 5B**). On the other hand, PC2 is more correlated with % fungal load and DON concentration, indicating that this PC is associated with severity of infection (**Fig. 5B**). Genes that contributed to PC1 and PC2 are highlighted in **Fig. 5C**. A more careful examination of the PCA plots reveals that plants that have high values of PC1 and PC2 tend to have high AUDPC (**Fig. 5D**) and DON (**Fig. 5E**). On the other hand, plants that had low PC1 and PC2 tended to have high AUDPC and percent fungal RNA (**Fig. 5F**). This reveals that plants containing genetic variants that have low PC1 and PC2 would have an optimal combination of traits. The genetic variants found in the GWAS analysis were associated with fusarium symptoms in an independent set of plants, demonstrating the potential of our approach for suggesting targets for marker-assisted selection of Fusarium resistance.

## Discussion

We set out to characterise symptom heterogeneity within a single barley cultivar, AAC Synergy, with the aims of (i) associating early gene expression with later disease-related traits (ii) identifying variants that are already present within this elite cultivar that could be used in marker-assisted selection of more *Fusarium* resistant barley plants. Our work provided fundamental insights into early response to *F. graminearum* infection in barley. In doing so, we demonstrate that single plant-omics is a powerful new tool for investigating plant-pathogen interactions.

Our single plant transcriptomics approach yielded critical insights into the relationship between fungal load, visible symptoms and DON mycotoxins within the barley population. Early fungal load was not associated with later symptoms. Gene expression and % fungal RNA measurements are derived from the top two spikelets, while AUDPC and DON information is derived from the remainder of the head. Therefore, the weak association between percent fungal RNA and later symptoms may be a result of spatial heterogeneity in fungal load. An alternative explanation is that heterogeneity in immune responses within the population weakens the association between initial fungal load and later symptoms.

Genes that were differentially expressed early in *F. graminearum* infection were associated with the current fungal load and with later visible symptoms (AUDPC) and DON mycotoxin concentrations. Genes that were up-regulated during infection were more closely associated with fungal RNA, while genes that were down-regulated during infection were more closely associated with later symptom severity. What is unclear is whether the early downregulation of this gene set caused increased severity of symptoms or whether observed downregulation is an indicator that the plant’s immune response was not effectively coping with the infection.

Overall, our approach was able to uncover most major known pathways involved in response to *F. graminearum* challenge (**Fig. 4**), highlighting several key players associated with visible symptoms and DON mycotoxin concentrations.

The gene families associated with lowered AUDPC included those predicted to encode protein kinases (CDPKs or CPKs)^29,30^, plant lectins (ricin B-like lectin)^31^, actin cytoskeleton^32,33^, reactive oxygen species (ROS) activity^28^, E3 ubiquitin ligases^34^ and zinc finger transcription factors^35,36^. These families have a well-studied role in defence response, playing an integral role in extracellular recognition of pathogens and producing an immune response (often collectively referred to as PTI). It is hard to assess whether upregulation of receptor-kinases and similar plant immunity proteins is a response to pathogen infection or causes a reduction in symptoms, or if successful infection reduces the expression of critical R-genes. However, conventional wisdom regarding plant immunity indicates that the latter is more likely: in general R-genes are found to be highly expressed in steady-state growth conditions (eg in sufficient quantity that their stimulation produces an immediate and effective reaction). It is unclear how a recognition event producing increased receptor transcription and translation would provide a short-term benefit to the plant. Instead, the recognition event would be expected to have a regulatory effect on the downstream mechanisms of defence such as via zinc-finger transcription factors and subsequent synthesis of antimicrobials, cell wall strengthening, withdrawal of resources from the infected area, etc. Furthermore, it is well established that successful biotrophic or hemi-biotrophic infection suppresses the plant immune response against all pathogens, not just the infecting species, as secondary infection by an avirulent pathovar is possible once the immune system has been suppressed by the virulent pathovar^37^.

The genes associated with DON mycotoxin levels were primarily glycosyltransferases/beta-glucosidases. This category of protein has been previously identified as having a role in detoxification of DON via glycosylation and preventing its effects^25,38^. Additional genes associated with low DON levels and not with low infection in general were those associated with lipid and polysaccharide metabolism, and cell cycle/homeostasis (including ATP synthase)^39,40^. In this case, the role of these genes is unclear: it may be that high levels of DON disrupts these pathways to the pathogen’s benefit rather than that their expression produces low DON levels.

A central goal in studying plant-pathogen interactions is to understand the sequence of regulatory processes that occur during the onset of infection. The most common experimental approach for investigating these dynamics is a time-series design, in which samples are collected at regular intervals following infection. However, individual plants often progress through infection at different rates due to asynchronous responses, making fine-scale mapping of regulatory processes difficult. An alternative strategy is to leverage this asynchrony. By sampling individual plants that were infected simultaneously but are at different stages of infection, we can order them according to their degree of disease progression (a pseudotime). This allows us to observe how gene expression patterns change along the pseudotime axis, providing a clearer picture of the transcriptional sequence underlying early infection. One limitation of this approach is that, while it can reveal overarching trajectories of transcriptional states across plants, it does not identify the source of heterogeneity. For example, differences among trajectories could arise from variation in the initial fungal load, differences in the rate of immune response, or a combination of both factors.

Among genes that had higher expression under *Fg*-treated barley than controls, we found one cluster that was expressed early in the pseudotime that included chitinase activity and calcium binding. These processes are both important in early infection, where calcium ion fluxes act as early intracellular signals in response to fungal elicitors like chitin oligomers triggering downstream immune response and chitinases physically break chitins in fungal cell walls or cause cell wall remodelling during infection ^41,42^. Glutothionine-related genes are expressed next in our pseudotime. Glutothionine is known to modulate hormone and redox responses to infection, as well as trigger leaf-to-leaf calcium signalling^43^. This phase also coincides with the activation of glycosylotransferase genes, which play a pivotal role in DON detoxification, as previously highlighted^25^. Finally, late in the pseudotime genes are expressed that are associated with the extracellular region and strictosidine synthase, which catalyses the first step of monoterpene alkaloid synthesis, a key aspect of a plants’ chemical defense.

This pseudotime was strongly associated with percent fungal RNA and, to a lesser extent, later symptoms. Therefore, the inherent transcriptional trajectory within the population is biologically meaningful, although we cannot determine whether initial fungal load or heterogeneous responses to infection led to these trajectories.

A key outcome of the analysis is a collection of genetic variants that are associated with transcriptional trajectories (pseudotimes), percent fungal RNA, visible symptoms and DON concentrations. Almost every one of these variants are within known defense-related genes, suggesting that these variants are enriched for causal variants. This is in contrast with traditional GWAS outcomes, where the causal variant is often hidden within a large linkage disequilibrium block^17^. The variants include those within a respiratory burst oxidase B-like, which generates oxidative bursts early in the fungal defense response^45^ and COP9 signalosome, a key ubiquitination factor that orchestrates ROS and hormone signalling in response to pathogens^46^.

The collection of variants that were identified are significantly associated with disease symptom severity in an independent set of plants, suggesting that these variants could be used in marker-assisted selection to develop sub-populations of AAC Synergy plants with reduced Fusarium susceptibility. Since these genetic variants all exist endemically within the AAC Synergy population, a high-yielding variety, and are only weakly associated with traits like dry mass, they are less likely to have a pronounced negative impact on yield-related traits than a Fusarium-resistance variant that is identified from a non-elite variety.

Single plant-omics has been used extensively to investigate plant developmental transitions, but here we show that it has much broader applications in investigating other traits with asynchronous timing, such as defense.

The approach taken here recapitulates a good deal of the past 20 years of molecular research into plant immune systems (**Fig. 4**). Without deliberately setting out to identify known components of the conventional plant defense response pathways, our single plant omics approach highlighted several of the main pathways associated with a complex, multi-phenotypic pathogen. The seemingly homogenous population of barley, grown in highly controlled growth chambers experienced sufficient variation in gene expression due to genetic, environmental, temporal, and stochiometric conditions, enabling us to identify transcriptional markers and genetic markers for *F. graminearum* disease symptoms. Given the difficulty of identifying QTLs for resistance breeding in FHB, thanks to its complex phenotype and high variability in virulence in experimental conditions which would produce easily measurable and consistent outcomes in a more tractable system, we propose that this approach is an invaluable supplement to traditional genome-based approaches to mapping disease resistance. In particular, the identification of LecRK-A4.3 and PERK9 as genes in AAC Synergy which are involved in a dosage-dependent manner with *F. graminearum* response permits empirical testing of these genes and subsequent integration of high-dosage variants into barley germplasm, as well as investigation of orthologous genes in other critical species such as bread wheat.

## Author contributions

EJR, SH, GSB, and DE conceived of the project.

ML, MJAA, YZ, JG, JV, CH, and DO contributed to the experimental work. EJR, SH, JH, BS, and DE contributed to the data analyses.

SH, ML, PD, RS, GSB, and DE contributed to the interpretation and contextualisation of the work.

EJR, SH, ML, GSB, and DE contributed to the writing of the manuscript. GSB and DE contributed to funding acquisition and project supervision. All authors contributed to the editing of the manuscript.

## Acknowledgements

This project was undertaken on the Viking Cluster, which is a high-performance compute facility provided by the University of York. The authors are grateful for computational support from the University of York high-performance computing service, Viking, and the Research Computing team. EJR discloses support from Biotechnology and Biological Sciences Research Council (BBSRC) – White Rose DTP in Mechanistic Biology [grant number BB/T007222/1, Ref.: 2444228]. DE discloses support for the research of this work from BBSRC [grant number BB/V006665/1]. GSB acknowledges funding from Western Grains Research Foundation (Project ID: AGR 2322), Saskatchewan Barley Development Commission (Project ID: 5116), and Natural Sciences and Engineering Research Council of Canada (NSERC) (Project ID: ALLRP 587124-2023).

## Online Methods

### Plant material and growth conditions

As well as being the most popular malting barley cultivar in western Canada, AAC Synergy (pedigree: TR02267/Newdale; TR02267 = T253/AC Metcalfe) is derived from elite Canadian barley breeding germplasm, and shares alleles with many other lines grown in the field. AAC Synergy is a Canadian 2-row malting spring barley cultivar developed in 2012 by Agriculture and Agrifood Canada^18^. Seeds (Breeder seed source) of barley cultivar AAC Synergy were obtained from Agriculture and Agri-Food Canada. Three plants were grown to maturity under experimental conditions and self-pollinated to generate the seed stock used in this study.

Seeds were sown individually into 1 L vertical tube pots (30 cm depth) filled with Sunshine Mix #1 (Sun Gro, Canada) supplemented with Miracle-Gro slow-release fertilizer. Plants were cultivated in a Conviron PGR15 reach-in growth chamber (Conviron, Winnipeg, Canada) under a 24/19°C day/night cycle, with a 16 h photoperiod, 800 lumens LED light intensity, and 70% relative humidity. Post-inoculation humidity occasionally rose to 80–90% but returned to baseline within one hour. Plants were watered every other day at the base to avoid wetting the heads and upper leaves. Weekly monitoring included measurements of height and developmental stage, and plants were rotated within the chamber to ensure uniform exposure to light and humidity. Inoculation with *Fusarium graminearum* spore suspension or distilled water procedural control was performed at anthesis, determined by manual inspection.

Treatment assignment was randomized using the Excel RAND() function. Physiological measurements are detailed in Supplementary Data Set 1.

### Fungal inoculum preparation

The *F. graminearum* strain SK-17-97, a 3ADON chemotype isolated from wheat in Saskatchewan, was used for inoculation. A five-day-old colony grown on 2% PDA at 24°C in darkness was fragmented and transferred to 200 mL of carboxymethyl cellulose (CMC) liquid medium (15% CMC, 1% yeast extract, 1% MgSO₄, 1% NH₄NO₃, 1% KH₂PO₄) supplemented with 0.2 g streptomycin. Cultures were incubated for five days with constant aeration at 24°C. The resulting spore suspension was filtered through Miracloth (Millipore), diluted to 6×10⁴ spores/mL in ddH₂O, and stored in 10 mL aliquots at −80°C.

### Disease inoculation and phenotyping

During half-anthesis (Zadoks stage 65), heads were spray-inoculated with 1 mL of conidial suspension or ddH₂O control from a distance of 10 cm, following removal of the flag leaf. Inoculated heads were enclosed in pre-wetted plastic bags for 48 hours to facilitate infection. Each plant received one inoculation, typically on the primary head, with additional heads treated when development permitted. Disease severity was assessed at 7-, 11-, 15-, and 20-days post-inoculation (dpi) by calculating the proportion of spikelets exhibiting white mycelium. The area under the disease progress curve (AUDPC) was computed using the formula: proportion × number of days^47^. Secondary infections and mechanical damage were recorded. Mature heads were retained for DON analysis. Inoculation and infection tracking details are provided in Supplementary Data Set 1.

### DON quantification and chemotype profiling

Mature infected and control heads were dried with silica gel for three days, weighed, and ground into fine powder using a Qiagen Tissuelyser II (Qiagen, USA). Samples were aliquoted into tubes containing <= 100 mg of tissue. Each aliquot was extracted with 1 mL of LC-MS grade acetonitrile containing 0.1% formic acid, nutated for 1 hour at room temperature, and centrifuged at 10,000 × g for 2 minutes. Extracts were analyzed using an LTQ Orbitrap XL Hybrid Ion Trap mass spectrometer (Thermo Scientific) coupled to a Dionex Ultimate 3000 UHPLC system. Separation was performed on a Kinetex C18 column (50 mm × 2.1 mm, 1.7 µm; Phenomenex) at 30°C with a flow rate of 0.35 mL/min. Mobile phases were water (A) and acetonitrile (B), both with 0.1% formic acid. The gradient included: 95% A for 0.5 min, linear ramp to 95% B over 4 min, hold at 95% B for 3.5 min, and re-equilibration at 95% A for 7 min.

HRMS was conducted in positive electrospray ionization mode (ESI⁺) with the following settings: m/z 100–2000, resolution 30,000; sheath gas 40; auxiliary gas 5; sweep gas 2; source voltage 4.0 kV; capillary temperature 320°C; capillary voltage 35 V; tube lens voltage 100 V. Samples were injected in randomized order. DON and 3/15ADON were quantified using in-house standards and Xcalibur 4.1 Quan Browser (Thermo Scientific), with result units expressed as µg per plant or µg per mg of tissue. Supplementary Data Set 1 includes tissue weights and DON concentrations.

### RNA extraction and sequencing

At 5 dpi, the top two spikelets (one floret/six kernels) were harvested for RNA extraction. Florets were excised using ethanol-sterilized scissors and flash-frozen in liquid nitrogen. Samples were ground in liquid nitrogen using a ceramic mortar and pestle, and RNA was extracted using the Qiagen RNeasy Mini Kit per manufacturer’s instructions. RNA integrity was assessed via 1% agarose gel electrophoresis stained with ethidium bromide, and concentration was measured using a Nanodrop spectrophotometer (Implen, USA). Samples with secondary infections or mechanical damage were excluded. High-quality RNA was sent to Novogene Corporation Inc. (USA) for library preparation and sequencing (60 million paired-end 150 bp reads) on the Illumina NovaSeq platform (Genomic Quebec, Canada). Sample metadata are linked in Supplementary Data Set 1.

### Gene prediction and annotation

Gene annotation was performed using the AAC Synergy v1.0 genome^48^ (Xu et al., 2021), downloaded from NCBI. Repetitive elements were annotated using RepeatMasker v4.1.2 with the Ensembl nrTEplantsJune2020 database (Smit et al., 2013–2015). The soft-masked genome was aligned to RNA-seq reads using STAR v2.7.10b^49^ with parameters: --twopassMode, --outFilterMismatchNmax 5, --outFilterMatchNminOverLread 0.80, --alignMatesGapMax 100000. The resulting BAM files and genome were input into the Braker3 pipeline for gene prediction, using the OrthoDB v11 Viridiplantae proteome^50,51^. Gene models were annotated using InterProScan v5.64-96.0^52^. Post-processing (e.g., GFF3 merging, conversion to.fna/.faa) was performed using the AGAT toolkit^53^.

### Genetic variant calling Data filtering steps

Only genes whose sum of their CPM across all samples was greater than one were used in our analysis, with the threshold determined by visual inspection of an elbow plot (Fig S1A). A Principal Component Analysis (PCA) was performed (prcomp, *stats* package version 4.3.2) on the variants (0 is homozygous matching reference genome, 1 heterozygous, 2 is homozygous different from reference). Two distinct clusters were identified (Fig S1B) across the first principal component (PC), and all variants with an absolute value of the loadings greater than 0.02 along this first PC were removed from the subsequent analysis. Furthermore, genes that were differentially expressed between these two clusters were identified by a non-parametric Mann-Whitney U test^54^. All genes with a p-value less than 0.05 and whose absolute value of the log2 fold-change in expression between the two groups was greater than 1 were also filtered out from future transcriptomic analysis.

### Differential expression analysis

Three different differentially expression analyses were performed: (i) Fusarium treatment (n=121) versus control group (n=41) (ii) top 25% versus bottom 25% AUDPC (within Fusarium treatment group) (iii) top 25% versus bottom 25% total DON µg/mg (within Fusarium treatment group). Comparison (i) was performed using a nonparametric Mann-Whitney U-test^54^, due to the large number of replicates, while (ii) and (iii) were performed using DESeq2 (version1.48.2)^55^, using the raw read counts. In all cases, the p-value threshold was set at 0.05 and the log2-fold-change threshold was set at 1. For each gene that was identified as being differentially expressed, a Spearman’s correlation was calculated between its expression levels and the measured traits in each barley plant.

### GO term enrichment analysis

GO terms for each predicted protein were identified using InterProScan version 101.0^52^, and then genes were clustered according to each shared GO term using the enricher function of the clusterprofiler R package version 4.16.0, with Benjamini-Hochberg adjusted p-values^56^. The GOfuncR package was used to format the supplementary table^57^. Families with overlapping GO terms (e.g. iron binding, and heme binding) were manually evaluated at the end of the analysis rather than pre-processed. GO term enrichment for each gene set was constructed

### Pseudotime analysis

The pseudotime analysis was only performed on plants that were treated with Fusarium and where only a single head was sampled. Two different gene sets were used to generate pseudotimes: genes with higher expression under treatment (up set) and genes with lower expression under treatment (down set). In both cases, we partitioned these gene lists into 300 equal sets of genes. For each partition, we used the pseudotime inference method from Redmond et al. (2024)^13^. First, we scaled the gene expression values of each to be between 0 (minimum expression) and 1 (maximum expression), then took a sum of those genes’ expression for each plant. Next, we ranked the plants from lowest to highest expression. We observed that the pseudotimes were consistent across the partitions so we determined the final pseudotime by ordering the plants by their average rank. The outcome of this method is that– overall– genes within this set will increase in their expression over the pseudotime. Gene expression over pseudotime was visualised using the pheatmap package version 1.0.13, with clusters determined by complete linkage hierarchical clustering^58^. GO enrichment for each cluster was determined, as previously described. For each pseudotime (generated from the up and down gene sets), Spearman’s correlation was calculated between the pseudotime rank and the traits measured.

### Genome-wide association study

There are not many genetic variants within gene bodies within a single cultivar of barley, so we hypothesised that some of these are likely causally linked to phenotypes. For this reason, we only performed a very limited filtering of the variant list to construct a genetic map for GWAS. First, variants were filtered to only include ones where the major genotype was found in less than 90% of the samples, leaving 3198 variants in the genetic map. The statgenGWAS package was used to perform a GWAS analysis^59^, with kinship calculated using the Identical by State method and significance calculated by an False Discovery Rate metric (p-value threshold of 0.05) as proposed by Brzyski D.^60^ which identifies phenotype-aware representatives from each cluster of co-localised markers– a strategy that seemed most aligned with our decision to only do light filtering of the genetic map. Only plants that were treated with fusarium and where only one sample was sequenced by RNA-seq were included in this analysis. Manhattan plots were generated. To further investigate the significant variants, the 400bp region around the variants were extracted from the genomic DNA and then searched in NCBI BLASTx version 2.17.0 (clustered nr database:https://blast.ncbi.nlm.nih.gov/Blast.cgi) to identify the closest orthologs^61^.

### Validation of markers

Our validation set consisted of plants that were treated with fusarium and had two heads separately sampled. All genetic variants that were associated with pseudotime up, pseudotime down, AUDPC, Total DON ug per mg, or percent fungal RNA were compiled. These variants were used to perform a PCA, with scaling prior to eigenvector decomposition, using the prcomp package^62^. Spearman correlation was calculated between the two first principal components and the average value of the traits between the two technical replicates.

