## Supplemental Figures for "Harnessing within-cultivar variation to identify hidden genetic resistance using single plant-omics"

### Supplementary figures to: “Harnessing within-cultivar variation to predict genetic resistance using single plant-omics”

Ethan J Redmond*[1], Meng Li[2]*, Samuel Holden[2]*, Muhammad Jawad Akbar Awan[2], Yishan Zhang[3], Jujhar Singh Gill[3], Jashanpreet Virhia[2], Jess Hargreaves[4], Peter Danks[1], Beth Sleath[1], Rajagopal Subramaniam[5], Carmen Hicks[5], David Overy[5] Gurcharn Singh Brar[2][3]*, Daphne Ezer[1]*

[1] Department of Biology, University of York, Heslington, York United Kingdom, YO10 5DD.

[2] Faculty of Agricultural, Land and Environmental Systems, University of Alberta, Edmonton, AB, Canada, T6G 2R3.

[3] Faculty of Land and Food Systems, The University of British Columbia, Vancouver, BC, Canada, V6T 1Z4

[4] Department of Mathematics, University of York, Heslington, York United Kingdom, YO10 5DD

[5] Agriculture and Agrifood Canada Ottawa Research and Development Centre, Ottawa, ON, Canada, K1A 0C6

**Description of Supplementary Tables**

**Table S1:** Phenotype tables for fusarium and mock-treated AAC Synergy plants, including AUDPC and DON metrics.

**Table S2**: RNA-seq data for fusarium and mock-treated AAC Synergy plants and within-gene body genetic variants. A look-up table for gene names to nearest homolog is provided to contextualise these results.

**Table S3**: Differential gene expression sets (two sub-populations; treatment v control; high AUDPC vs low AUDPC)

**Table S4:** GO terms within gene clusters of differentially expressed gene sets, and corresponding gene sets. This table includes GO terms associated with pseudotime clusters.

**Table S5:** GWAS outcomes and BLASTx hits for genetic variants with significant p values.

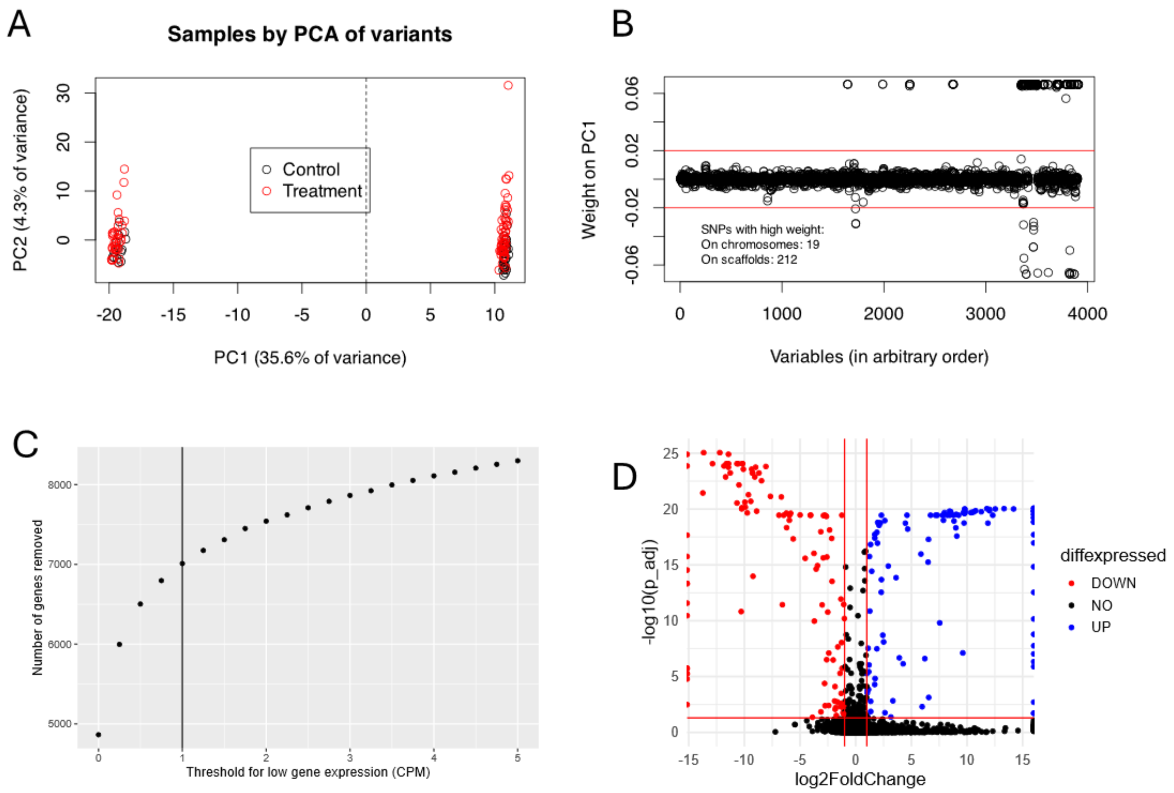


**Figure S1: Motivation for filtering genes in further analysis**. (A) A PCA of the variants suggested that the plants formed two separate clusters along PC1. (B) Of the variants that have high weights along PC1 (|weight|>0.02), only 19 mapped to chromosomes, while 212 mapped to scaffolds. Therefore, we chose not to include any genes that did not map to chromosomes in our further analysis to simplify our interpretation. (C) Genes whose counts per million were less than 1 across all RNA-seq samples were removed, as determined by an elbow-based approach. (D) Moreover, we decided to filter for any genes that were differentially expressed between the two sub-groups in our population, as indicated in red and blue in our volcano plot.

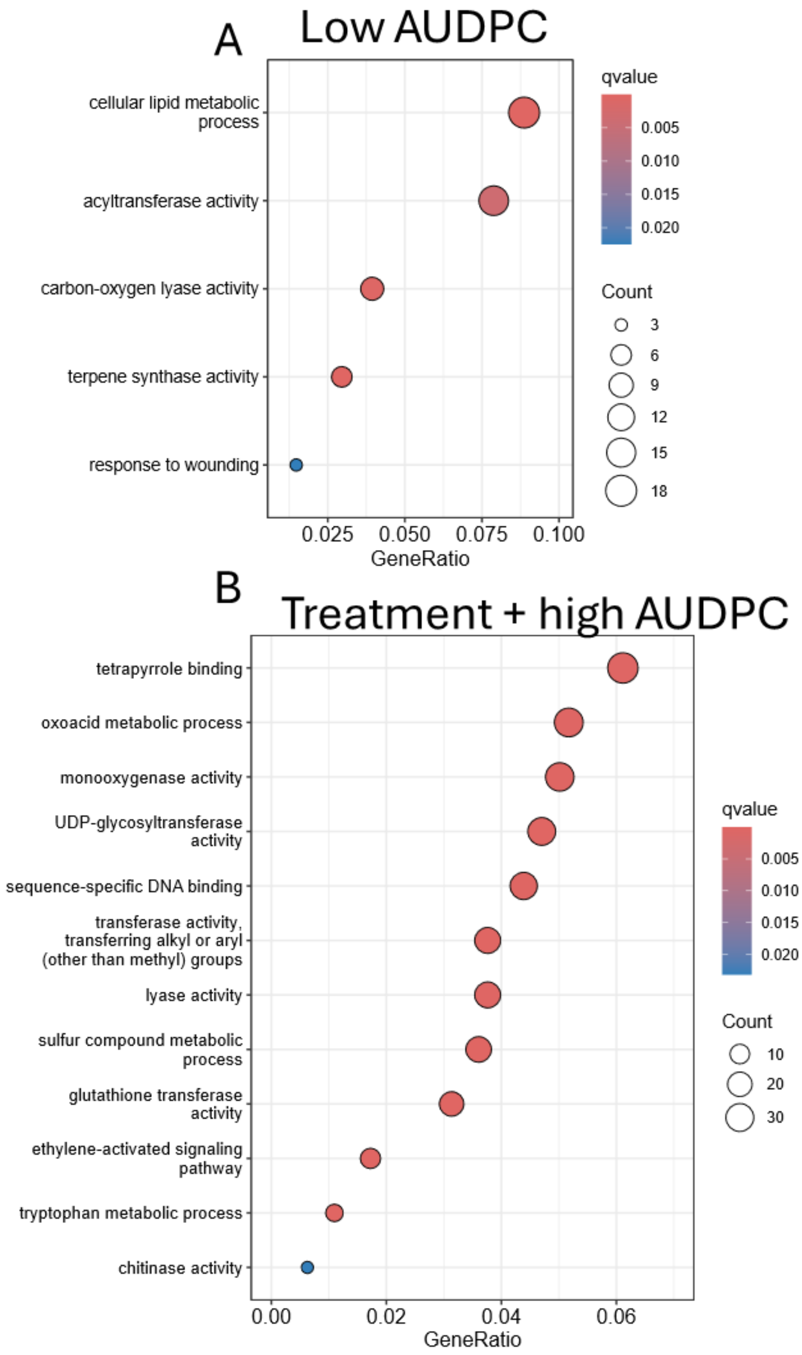


**Figure S2: GO term enrichment for additional gene sets.** Panel (A) shows GO term enrichment for genes that were more highly expressed in the bottom 25% AUDPC than the top 25% AUDPC. Panel (B) shows GO term enrichment for genes that were more highly expressed both in fusarium treated plants (vs mock-treated) and in the top 25% AUDPC (vs bottom 25% AUDPC).

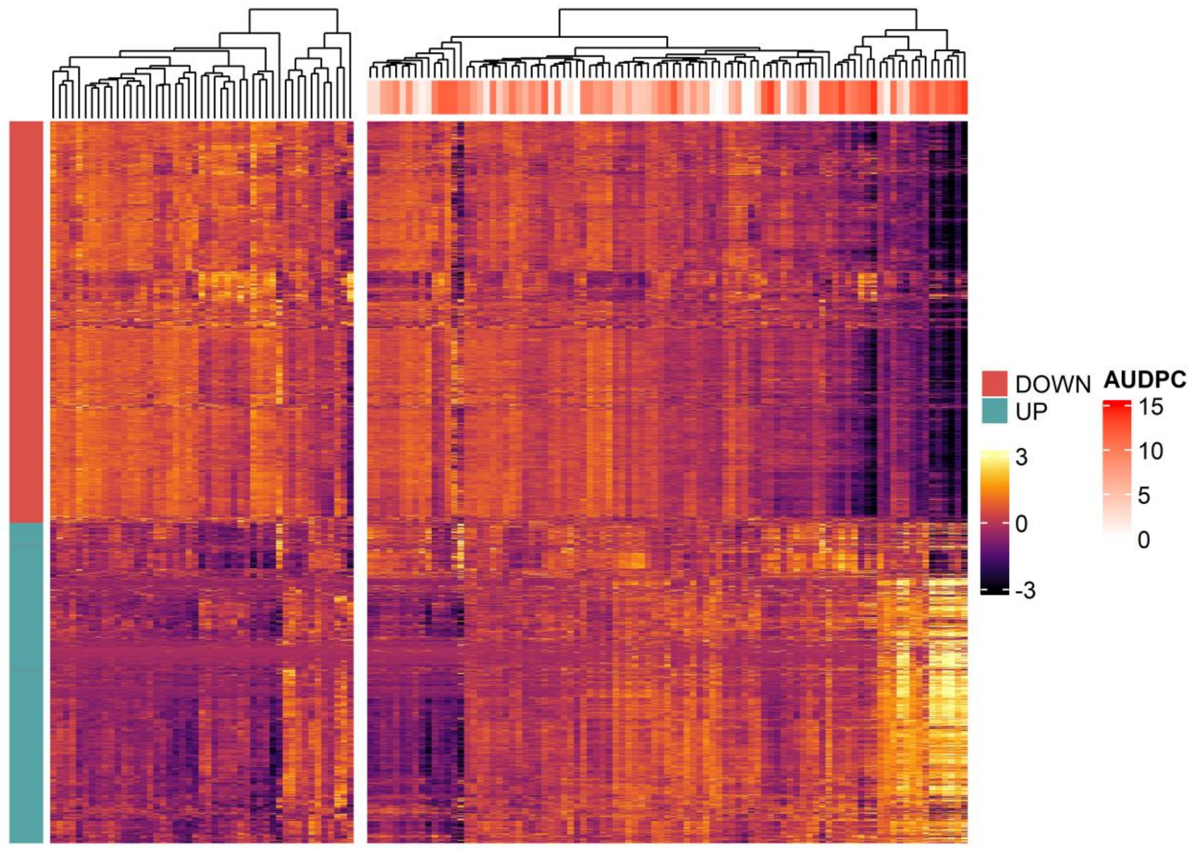


**Figure S3: Heatmap of differentially expressed genes (Treatment vs Control)**. The Z-score of the log-CPMs are shown here for all genes that were differentially expressed under fusarium treatment and control plants (rows). Each column represents an RNA-seq sample. Annotations of the rows show whether genes had increased (up) or reduced (down) expression under the fusarium treatment. Annotations of the column indicate AUCDP, a marker of infection severity. Many of the genes that had greater expression in the treatment are particularly highly expressed in a small group of high AUDPC samples and vice versa.

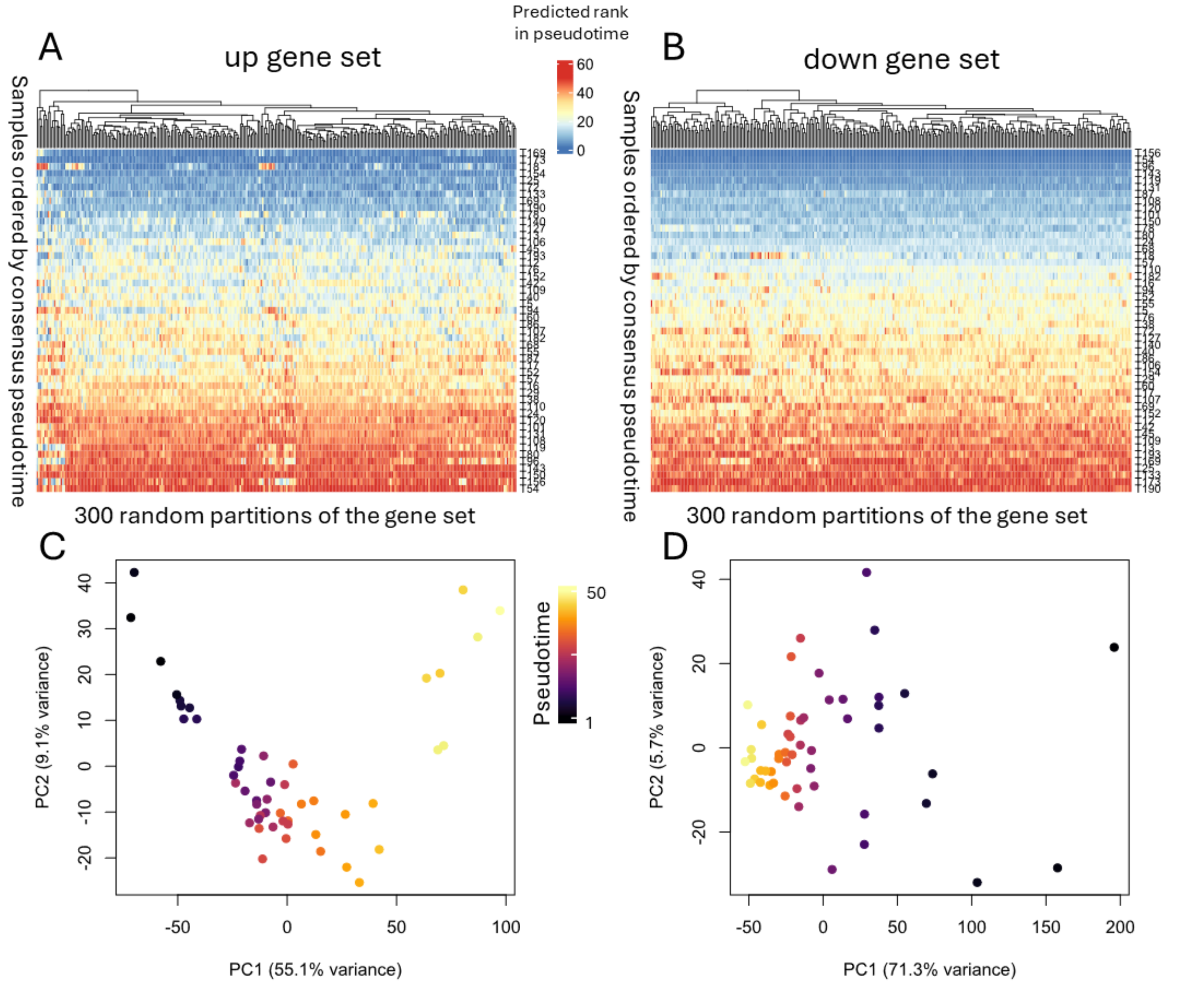


**Figure S4: Pseudotime inference**. Differentially expressed genes between control and treatment groups were used to construct a pseudotime. Each gene set was randomly partitioned into 300 equal groups, a sum of the scaled log-CPMs was taken, and the samples' rank corresponds to its pseudotime. The rank of the average pseudotime position was used as the consensus pseudotime. The pseudotimes were consistent across the 300 random partitions for both the genes whose expression was higher under fusarium infection – "up gene set" (A) and those whose expressions was lower under fusarium infection – "down gene set" (B). Columns were clustered using complete linkage and rows were ordered by the consensus pseudotime of the corresponding gene set. Principal Component Analysis (PCA) was performed on the scaled log-CPM tables, using either the up gene set (C) or down gene set (D). Colours indicate the position of the same in the consensus pseudotime. (C) shows a trajectory that aligns with the pseudotime, while in (D) we see that the pseudotime is associated with PC1, which explains 71.3% of the variance.

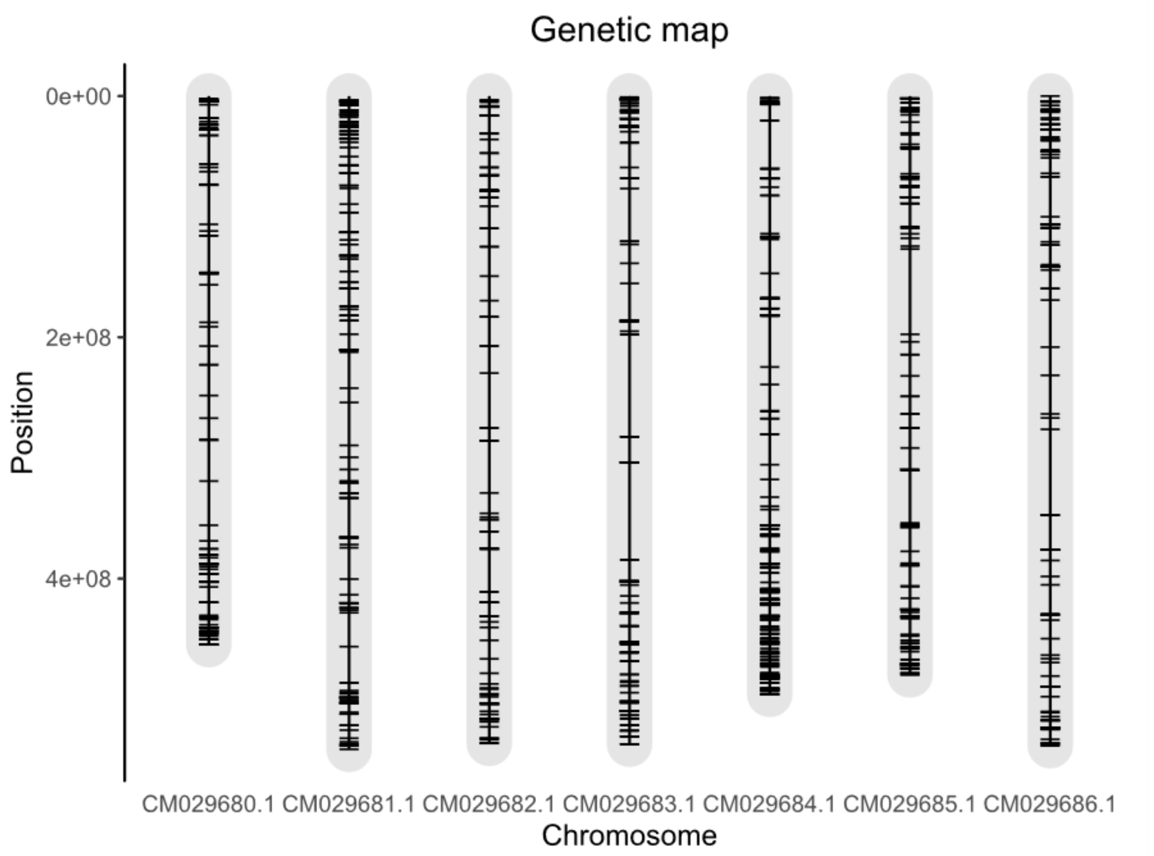


**Figure S5: Genetic map showing genomic locations of every high confidence variant found in more than 10% of plants.**

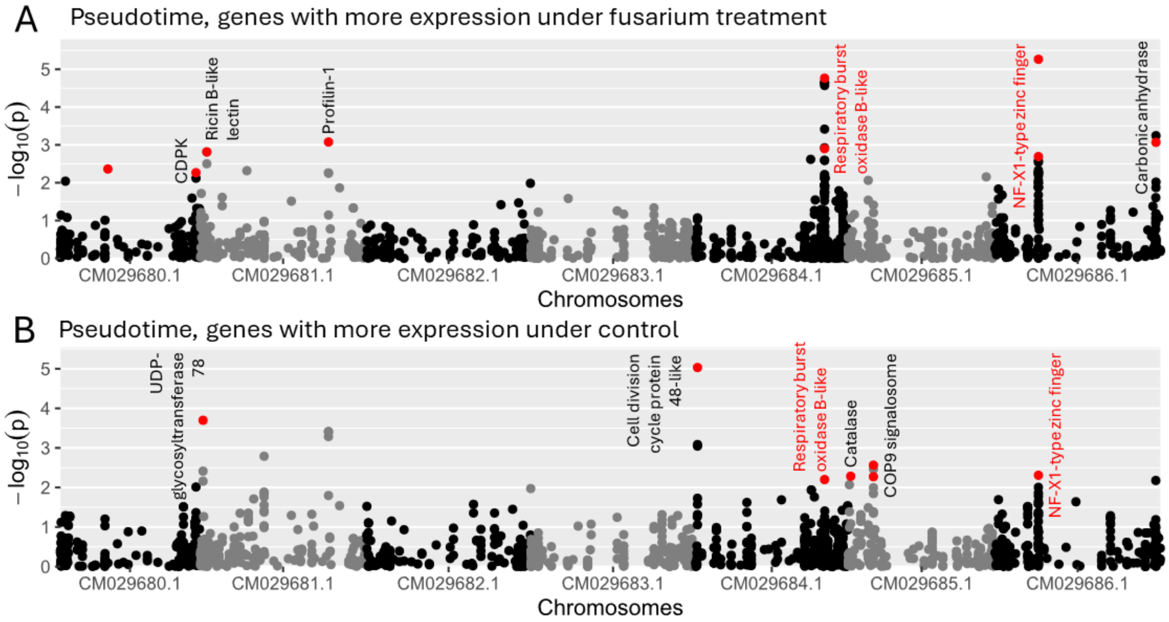


**Figure S6: Manhattan plots showing variants associated with pseudotimes.** Variants in red were identified based on a false discovery rate-based method, with the genes containing the variants written for reference. Genome-wide association was determined either using the pseudotime using genes that increase their expression under fusarium treatment (A) and those with more expression in controls (B). Variants that are found associated with both pseudotimes are highlighted in red.

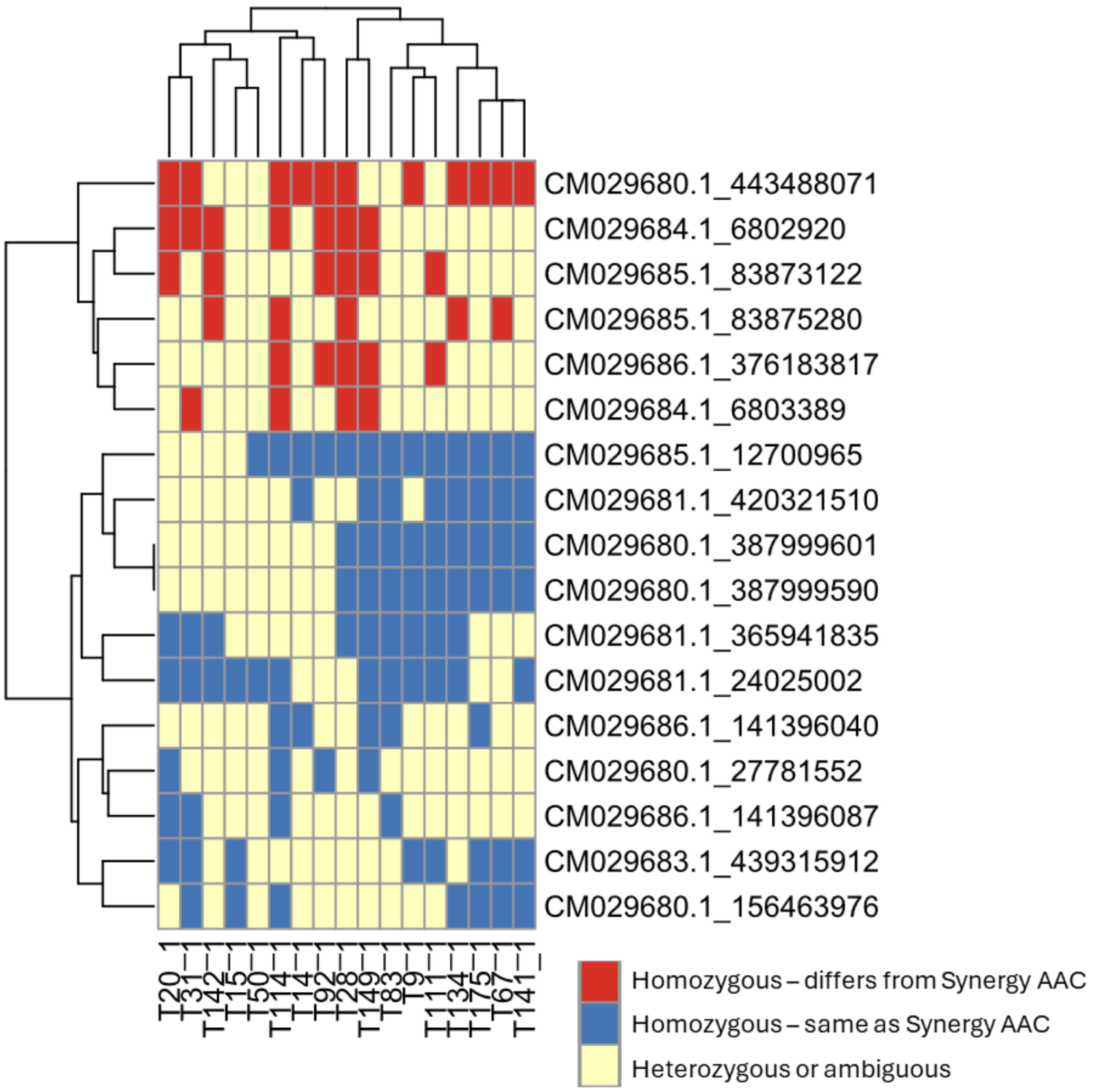


**Figure S7: Variants that were significant in at least one GWAS and that were found in the validation set are shown clustered here.** Rows show the variants, labelled by chromosome and position, while columns show the plants in the validation set. Colours indicate genotype.
